# Inhibition of PI3K p110δ activity reduces IgE production in IL-4 and anti-CD40 stimulated human B cell cultures

**DOI:** 10.1101/2022.08.09.503316

**Authors:** Anna Cutrina-Pons, Aloka De Sa, David J Fear, Hannah J Gould, Faruk Ramadani

**Author notes:** **Correspondence:** Faruk Ramadani, Randall Centre of Cell and Molecular Biophysics, King’s College London, London, SE1 1UL, UK. The first two authors contributed equally to this article, and both should be considered first author.

## Abstract

Phosphoinositide 3-kinase (PI3K) p110δ signalling negatively regulates the production of mouse IgE. However, there are disparities between the mouse and human IgE biology, and the role of PI3K p110δ in the production of human IgE is yet to be determined. To investigate the effect of PI3K p110δ inhibition in the production of human IgE we isolated human B cells from tonsil tissue and stimulated them with IL-4 and anti-CD40 antibody to induce class switching to IgE and IgG1 in the presence or absence of IC87114, a small molecule inhibitor of PI3K p110δ. Using FACS, RT-PCR and ELISA we examined the effect of PI3K p110δ inhibition on IgE production and also determined the mechanisms involved. Unlike in mice, we observed that PI3K p110δ inhibition significantly reduces the number of IgE+ switched cells and the amounts of secreted IgE in IL4 and anti-CD40 cultures. However, the number of IgG1+ cells and secreted IgG1 were largely unaffected by PI3K p110δ inhibition. The expression levels of AID, ε and γ1 germinal transcripts or other factors involved in the regulation of CSR to IgE and IgG1 were also unaffected by IC87114. However, we found that IC87114 significantly decreases the proliferation of tonsil B cells stimulated with IL-4 and anti-CD40, specifically reducing the frequency of cells that had undergone 4 divisions or more. Our data identifies proliferation as the key mechanism through which PI3K p110δ regulates CSR of human B cells to IgE and highlights PI3K p110δ as a pharmacological target for the treatment of allergic disease.

## Introduction

Immunoglobulin (IgE) antibodies play a fundamental role in the pathogenies of allergic disease. They bind to the high affinity IgE receptor (FcεRI) on mast cells and basophils, which, when cross-linked by multivalent allergens, trigger cell activation and the release of various mediators of immediate hypersensitivity and allergic inflammation (1,2). Since its discovery, over a half a century ago, we have made significant progress not only in understanding the immunological mechanisms associated with type 1 hypersensitivity reactions(1,2) but also in our understanding of the mechanisms involved in the regulation of IgE production(3–9).

The commitment of a B cell to undergo CSR to IgE is dependent on signals downstream of the tumour necrosis factor receptor superfamily member CD40, after its interaction with the CD40 ligand (CD40L) on the surface of T cells, and IL4/L13 binding to their receptors on B cells(2,10). These signals activate the nuclear factor-κ B (NFκB) and signal transducer and activator of transcription 6 (STAT6) transcription factors that act synergistically to regulate CSR to IgE(10–12). Both NFκB and STAT6 bind to the Iε promoter and initiate the synthesis of ε germline transcripts (εGLT) and make ε switch region available for genetic recombination. Furthermore, NFκB and STAT6 can also activate the expression of the enzyme activation-induced cytidine deaminase (AID)(13), an important regulator of CSR and SHM, which produces DNA breaks in the switch regions that precede the Cε exon(10).

Several molecular mechanisms have evolved to ensure rigorous regulation of IgE production(10,14). In mice, they also include signalling by PI3K p110δ(15–17), a member of the class IA PI3K catalytic isoforms, which is activated downstream of various receptors on B cells(18). Inhibition of the PI3K p110δ signalling in mouse B cells, both *in vitro* and *in vivo*, increases IgE production(15–17). This effect of PI3K p110δ inhibition was found to be associated with increased εGLT and AID expression levels, suggesting that under normal conditions PI3K p110δ signalling supresses IgE production by repressing εGLT and AID expression. This was surprising since signalling by PI3K p110δ is also required for the production of type 2 cytokines such as IL4 and IL13(19,20). It was later suggested that PI3K p110δ signalling limits IgE production in a B cell-intrinsic manner, a mechanism that involves the activation of Bcl6(16), a known negative regulator of CSR to IgE(21).

However, the B cell regulatory systems of IgE production in humans differ from those in mice(7), and the role of PI3K p110δ signalling in the production of human IgE is yet to be determined. Here, we isolated human CD43^-^ tonsil B cells and stimulated CSR to IgE with IL4 and anti-CD40. To investigate the involvement of PI3K p110δ signalling in human IgE production, B cell cultures were treated with IC87114, a small molecular inhibitor of p110δ, which was used to determine the role of PI3K p110δ signalling in IgE production by mouse B cells(15,17), and nemiralisib, a new generation of PI3K p110δ inhibitors (22). The data show that PI3K p110δ inhibition reduces IgE production in tonsil B cell cultures stimulated with IL-4 and anti-CD40 antibody. Interestingly, this effect was shown to be specific for IgE, with the production of IgG1 being largely unaffected. This result is in striking contrast with the effect of PI3K p110δ inhibition in mouse B cells(15–17,23).

## Materials and methods

### Ethics

Following full informed written consent, tonsils were obtained from patients undergoing routine tonsillectomies. The study was conducted at and in accordance with the recommendations of King’s College London and Guy’s and St Thomas’s NHS Fundation Trust and the protocol was approved by the London Bridge Research Ethics Committee (REC number 08/H0804/94).

### Cell culture

The untouched resting B cells were isolated from the human tonsils by depletion of other cells using MojoSort Human B Cell (CD43-) Isolation Kit (BioLegend), whereas the naïve B cells were isolated using the EasySep Human Naïve B cell isolation kit (Stem Cell Technologies). The purity of the B cells used in these experiments was > 95%. Purified B cells were then cultured at 0.5 ×10^6^ cells/mL in RPMI 1640 (Lonza) containing penicillin (100 IU/ml), streptomycin (100μg/ml) and glutamine (2mM, Invitrogen) and 10% Foetal Calf Serum (Hyclone; Perbio Biosciences). To induce CSR to IgE, cultures were stimulated with IL-4 (200 IU/ml; R&D Europe Systems Ltd) and anti-CD40 antibody (0.5 μg/ml; G28.5; American Type Culture Collection). To investigate the role of PI3K p110δ signalling in human IgE production, cell cultures were also treated with different concentration of IC87114 (LKT Laboratories Inc), a small molecule inhibitor of PI3K p110δ signalling used in the mouse studies. Considering that IC87114 has an IC50 of 0.5μM, it was used at concentrations below and above this IC50; 0.125μM, 0.25μM, 0.5μM, 1μM, and 2μM. PI3K p110δ was also inhibited using nemiralisb (GSK2269557), a highly selective inhibitor of p110δ (pKi=9.9) (MedChemExpress LLC). The cells were then incubated for up to 10 days at 37°C with 5% CO2.

### Flow cytometry

To determine the effect of the PI3K p110δ inhibition on STAT6 and NFκB phosphorylation levels, cells were harvested 24 hours after culture with IL-4 and anti-CD40 and fixed with 2% paraformaldehyde (PFA), washed with PBS and permeabilised with 1 ml of ice cold methanol for 20 mins on ice. The permeabilised cells were then intracellularly stained for 1 hour at room temperature (RT) with anti-STAT6Tyr641 Alexa Fluor 647 (A15137E; Biolegend) and anti-NFκB p65Ser536 Alexa Fluor 488 (93H1; Cell Signalling Technology). To determine the effect of the PI3K p110δ inhibition on CSR to IgE, cells were harvested on day 10 of culture and stained with a live/dead fixable stain dye (Life Technologies Ltd) followed by fixation with 2% paraformaldehyde and permeabilization with PBS containing 0.5% Triton X-100, and 0.5% saponin. The cells were then stained with anti-human IgE FITC (IgE21; eBioscience) and anti-IgG1-PE human (IS11-3B2.2.3; Miltenyi Biotec) for 30 minutes at room temperature in the dark. Following the staining cells were washed and resuspended in FACS buffer and acquired with BD FACS Canto.

### Proliferation

Following the CFSE labelling, performed using CellTrace™ CFSE Cell proliferation kit (C34554, Molecular Probes™ Invitrogen), the cells were cultured as above, with IL-4 and anti-CD40 antibody, in the presence or absence of 2µM IC87114. After 10 days of culture, cells were harvested and acquired using FACSCanto. The data was analysed using FlowJo and the proliferative capacity of cultured tonsil B cells was determined based on the dilution of the CFSE labelling.

### ELISA

Maxisorp plates (Nunc) were coated with polyclonal mouse anti-human IgE (Dako Agilent Technologies) or mouse anti-human IgG1 (BD Pharmingen) in pH 9.8 carbonate buffer overnight at 4°C. Unbound sites were blocked with 2.5% BSA (Sigma-Aldrich) for 2 hours at RT and then washed 4 times with PBS+0.05% Tween-20. Supernatants from day 10 cultures were then added at appropriate dilutions and plates incubated overnight at 4°C. Human serum IgE (NIBSC) or IgG1 (Sigma-Aldrich) were used to construct standard curves. Binding was detected by polyclonal goat anti-human IgE-HRP (Sigma-Aldrich) or mouse anti-human IgG HRP (BD Pharmingen) in 1% PBS/BSA for 2 h at 37°C. The color reaction was developed using the 1-StepTM Ultra TMB-ELISA substrate solution (Thermo Scientific) and plates were read at 450nm using the Labtech LT 4500 microplate reader. IgE and IgG1 concentrations were then calculated from the standard curve using Prism 9.2 software (GraphPad).

### qRT-PCR

Total RNA was isolated from cells using the Qiagen RNeasy Plus Mini Kit (Qiagen) and the RNA concentrations were measured using the NanoDrop 2000 (Thermo Scientific). The isolated RNA was then reverse transcribed into cDNA using SuperScript III First-Strand Synthesis SuperMix for qRT-PCR (Invitrogen). To evaluate the expression levels of certain genes we used TaqMan MGB gene expression assays and TaqMan Universal PCR Master Mix on Viia7 RT-PCR machine (Applied Biosystems). Gene expression was standardised to an endogenous reference gene 18s rRNA (Hs99999901_s1, Applied Biosystems). AID (Hs00757808_m1) and BCL6 (Hs00153368_m1) gene specific qPCR assays (Applied Biosystems) were used with Taqman MGB chemistry. εGLT and γ1GLT were analysed using previously published Taqman primers and probes (24). The sequences of forward and reverse primers for εGLT (fw: 5′-CTGTCCAGGAACCCGACAGA-3′; rev: 5′-TGCA GCAGCGGGTCAAG-3′) and γ1GLT (fw: 5’-CCAGGGCAGGGTCAGCA-3’; rev: 5’-GGTGCTCTTGGAGGAGGGT-3’) (Sigma-Aldrich) and their corresponding FAM-labelled MGB probes (εGLT: 5′-AGGCACCAAATG-3′ and γ1GLT: 5 ′-CTCAGCCAGGACCAAG-3′) (Applied Biosystems) were designed in house. All gene specific assays were multiplexed with the 18s endogenous control assay and run-in triplicate. SDS software was used to quantify the target cDNA relative to 18s according to the 2(-Δct) method.

## STATISTICAL ANALYSIS

Statistical analysis was performed using the One-Way ANOVA, with Tuky’s correction, or unless otherwise stated. A *p* value of < 0.05 was considered significant (\**p* < 0.05,\*\**p* < 0.01,\*\*\**p* < 0.001). Data shown represent mean +/-standard deviation (SD) or unless otherwise stated.

## RESULTS

### PI3K p110δ inhibition reduces CSR of human tonsil B cells to IgE *in vitro*

Studies in mice have shown that CSR to IgE can be enhanced both *in vivo* and *in vitro* when the activity of PI3K p110δ is inhibited(15–17). This effect of p110δ inhibition is not exclusive to CSR to IgE, with CSR to IgG1 also being enhanced in mouse B cells(15,17,23). To investigate whether CSR to IgE and IgG1 in human B cells is also affected by the inhibition of the p110δ signaling, as in the mouse system, the isolated untouched tonsil (CD43^-^) B cells were cultured with IL-4 and anti-CD40 and different concentrations of the IC87114. After 10 days of culture, we observed an IC87114 dose dependent reduction in the frequency of IgE^+^ cells (Figure 1A, B). While the number of IgE^+^ cells in cultures treated with 0.125µM and 0.25µM of IC87114 were not significantly different to the IL-4 and anti-CD40 cultures only, cultures treated with 0.5µM, 1µM and 2µM of IC87114 had a significant reduction in the number of IgE^+^ cells (Figure 1A, B). When quantifying the amounts of secreted IgE antibody in culture supernatants we observed that the effects of IC87114 on secreted IgE were much more drastic (Figure 1D). Indeed, cultures treated with 1µM and 2µM of IC87114 had almost 80% of their secreted IgE reduced compared to the IL-4 and anti-CD40 cultures only. Furthermore, we can also observe a significant reduction of the secreted IgE in cultures treated with 0.25µM of IC87114. Surprisingly, IC87114 had a negligible effect on the number of IgG1^+^ cells, and we could only observe a significant IgG1^+^ cell reduction in cultures treated with 2µM of IC87114 (Figure 1A, C). Similar effects of IC87114 on CSR were also observed in naïve B cell cultures stimulated with IL-4 and anti-CD40 (Figure 1F, G). The effects of IC87114 on IgE production were also confirmed using nemiralisb, one of the most selective PI3K p110δ currently in development (22). As can be seen in Figure 2, the percentages of IgE^+^ cells and the amounts of secreted IgE in cultures containing 1nM and 10nM nemiralisib are selectively reduced. These observations are in striking contrast to those reported by the mouse studies and suggests that PI3K p110δ regulation of CSR to IgE and IgG1 differs between the mouse and the human system.

**Figure 1.**
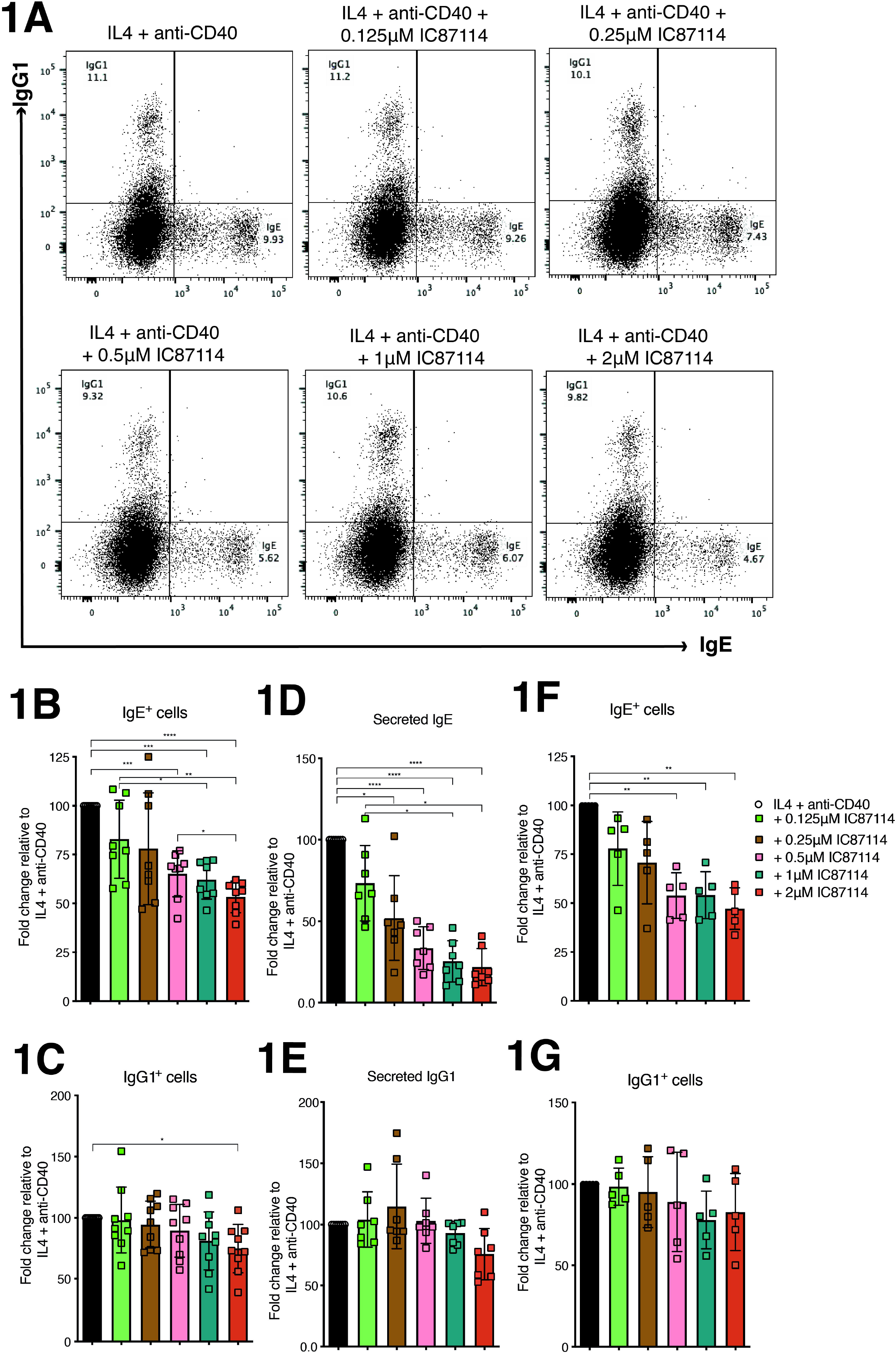
IC87114 reduces class switching of human B cells to IgE. (A) IL4 and anti-CD40 stimulated cells were harvested from the day 10 cultures containing different concentrations of IC87114 and intracellularly stained for IgE and IgG1. The percentages of total IgE^+^ cells (**B**) and IgG1^+^ cells (**C**) relative to the IL-4 and anti-CD40 only cultures. Secreted IgE (**D**) and IgG1(**E**) analyzed by ELISA. Data show the amounts of secreted IgE and IgG1 [ng/mL] made relative to the IL-4 and anti-CD40 only cultures. The percentages of total IgE^+^ cells (**F**) and IgG1^+^ cells (**G**) from naïve B cell cultures made relative to the IL-4 and anti-CD40 only cultures.

**Figure 2.**
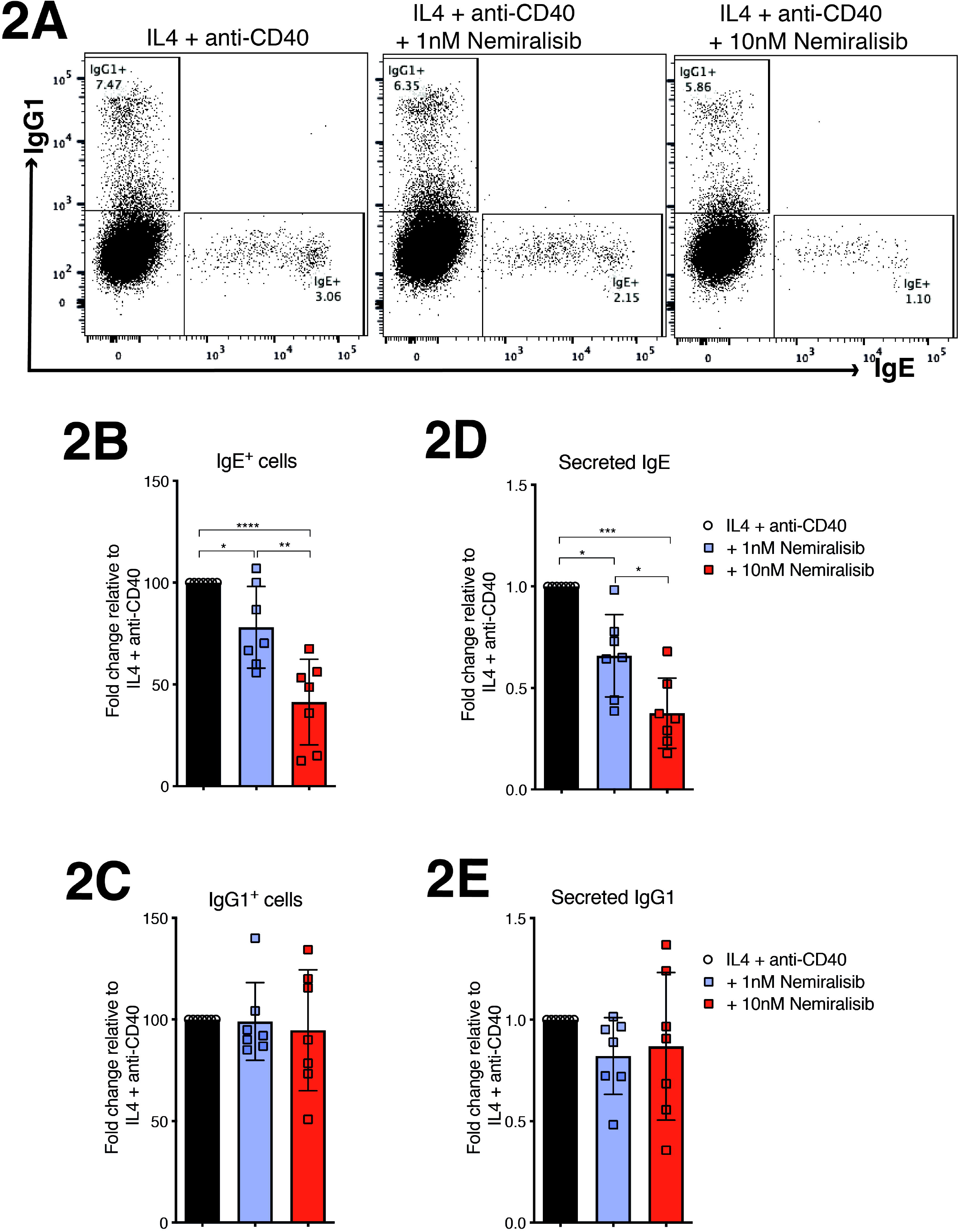
Inhibition of PI3K p110δ with nemiralisib selectively reduces class switching to IgE. Purified CD43-tonsil B cell cultures were stimulated with IL-4 and anti-CD40 alone or in the presence of 1nM and 10nM nemiralisib. (**A**) On day 10 of the culture, cells were harvested and intracellularly stained for IgE and IgG1. The percentages of total IgE^+^ cells (**B**) and IgG1^+^ cells (**C**) from several experiments were made relative to the IL-4 and anti-CD40 only cultures. Secreted IgE (**D**) and IgG1(**E**) analyzed by ELISA show the amounts of secreted IgE and IgG1 [ng/mL] made relative to the IL-4 and anti-CD40 only cultures.

### PI3K p110δ inhibition does not affect the class switch signals downstream of the IL-4R and CD40

To identify the mechanism by which PI3K p110δ regulates CSR to IgE and IgG1 in our cultured tonsil B cells, we first examined the expression levels of phosphorylated and thus activated, STAT6 and NFκB. As it can be seen in Figure 2A, 24 hours post culture with IL-4 and anti-CD40, the phosphorylation levels of STAT6 and NFκB were unaffected by the inhibition of p110δ PI3K with 2µM IC87114. However, previous mouse studies have reported that IC87114 regulates CSR to IgE and IgG1 by upregulating the expression of AID, εGLT and γ1GLT(15,17). Therefore, to confirm that the class switch signals downstream of the IL-4R and CD40 in human B cells are unaffected by the p110δ PI3K inhibition we also evaluated the expression levels of AID, εGLT and γ1GLT on day 5 of the cell culture. The RT-PCR data showed that the treatment of the cell culture with 2µM IC87114 has no significant effect in the expression levels of AID, εGLT and γ1GLT (Figure 2B). This is consistent with the unchanged STAT6 and NFκB phosphorylation levels and suggests that unlike in the mouse system, the effect of PI3K p110δ inhibition on CSR of human B cells to IgE and IgG1 occurs independently of class switch signals downstream of the IL-4R and CD40.

### PI3K p110δ inhibition does not affect the expression levels of BCL6 in human B cells

Previously, it was reported that PI3K p110δ regulates CSR of mouse B cells to IgE in a B cell intrinsic manner by modulating BCL6 expression(16). BCL6 is a transcription factor that is important for GC reaction but also is known to compete with STAT6 at the εGLT binding site leading to the repression of its transcription and that of CSR to IgE(10,21). Therefore, in mice the reduction of the BCL6 by the inhibition of p110δ can lead to enhanced CSR to IgE(16).

To test if this is also true in the human B cells, we evaluated the expression levels of BCL6 by RT-PCR. As can be seen from Figure 3, we did not observe any changes in the BCL6 expression levels between cell cultures that were untreated and those treated with 2µM IC87114.

**Figure 3.**
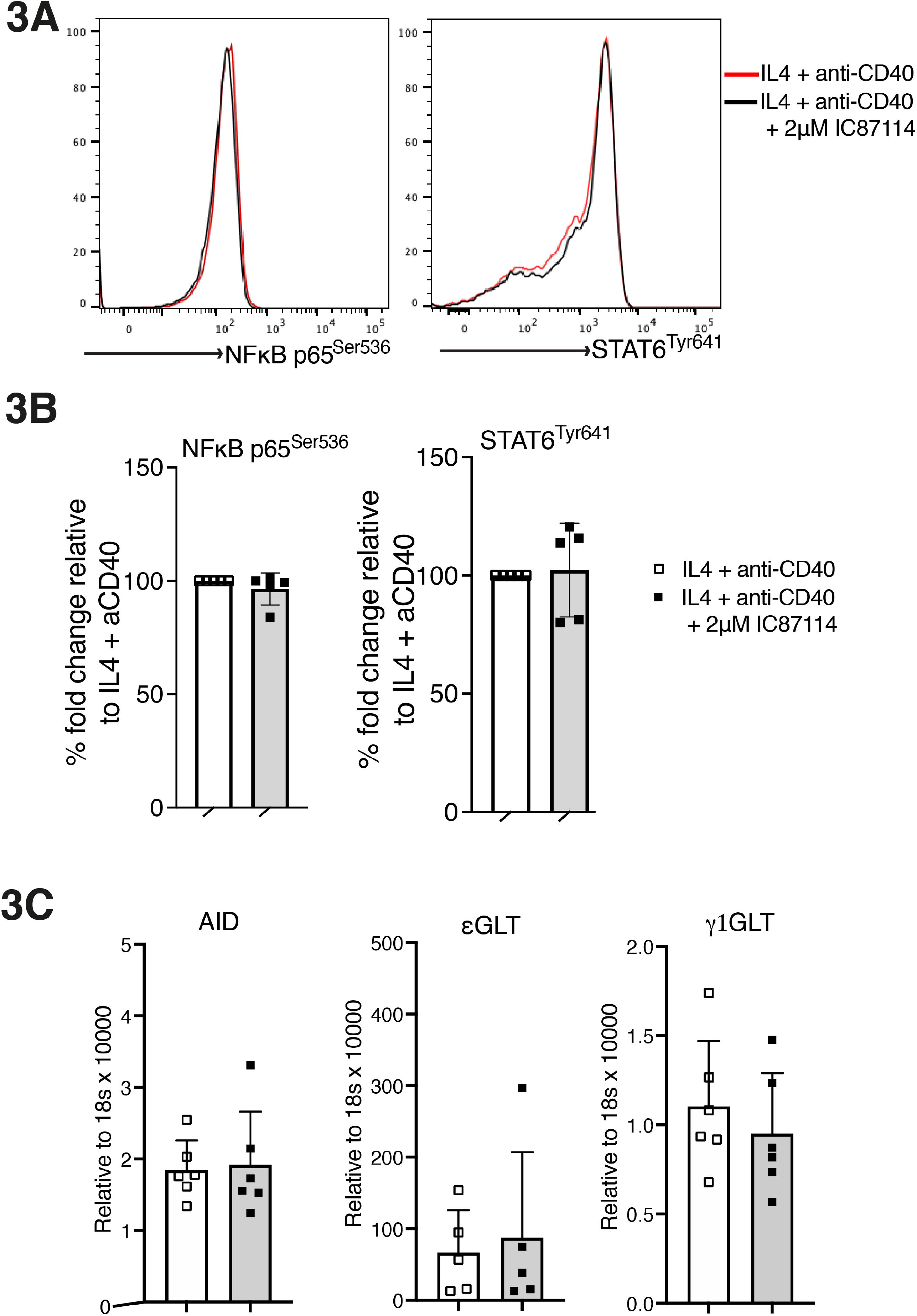
IC87114 has no effect on the class switch signalling downstream of IL-4R and CD40. After 24h of IL-4 and anti-CD40 stimulation, B cells were fixed, permeabilized and subsequently stained with anti-STAT6 phospho (Tyr641) and anti-NFκB p65 (Ser536). (**B**) The data show the fold change in median fluorescence intensity (MFI) relative to IL-4 and anti-CD40 only cultures. **(C)** The expressions of AID, εGLT and γ1GLT were quantified with qRT-PCR from RNA isolated from day 5 of the tonsillar B cell culture. Gene expression was normalised with 18s rRNA.

Overall, this data, and that from above, suggests that there are fundamental differences in the mechanisms through which PI3K p110δ signalling regulates CSR to IgE in human and mouse B cells.

### Inhibition of the PI3K p110δ activity reduces the proliferative capacity of human B cells stimulated to undergo CSR to IgE

Cell division plays an important role in CSR(25,26). For CSR to IgE to happen B cells are required to divide a minimum of 5 times, whereas CSR to IgG1 requires only 2 cell divisions(25,26). Mouse studies investigating the role of p110δ PI3K in IgE production did not report any significant effect of the p110δ PI3K inhibition on the IL-4 and anti-CD40 induced B cell proliferation(15,17,23). To determine if the p110δ PI3K inhibition reduces the proliferative capacity of the cultured tonsil B cells, thus, reducing the CSR of human B cells to IgE, we first labelled the isolated tonsil B cells with CFSE and analyzed cell division after 10 days of culture. The representative FACS histograms and cumulative data (Figure 4A, B) show that the number of proliferating B cells in cell cultures treated with 2µM IC87114 was reduced by half. Considering that the minimum number (5) of cell divisions required for CSR to IgE to occur, we also investigated the number of cell divisions by measuring the halving of the CFSE intensity. We found that the number of B cells undergoing 4 cell divisions, or more is significantly reduced when cell cultures were treated with 2µM IC87114 (Figure 4C, D). In contrast, the number of B cells undergoing 2 and 3 divisions, which are decisive for CSR to IgG1 (22) was slightly, but not significantly, reduced.

**Figure 4.**
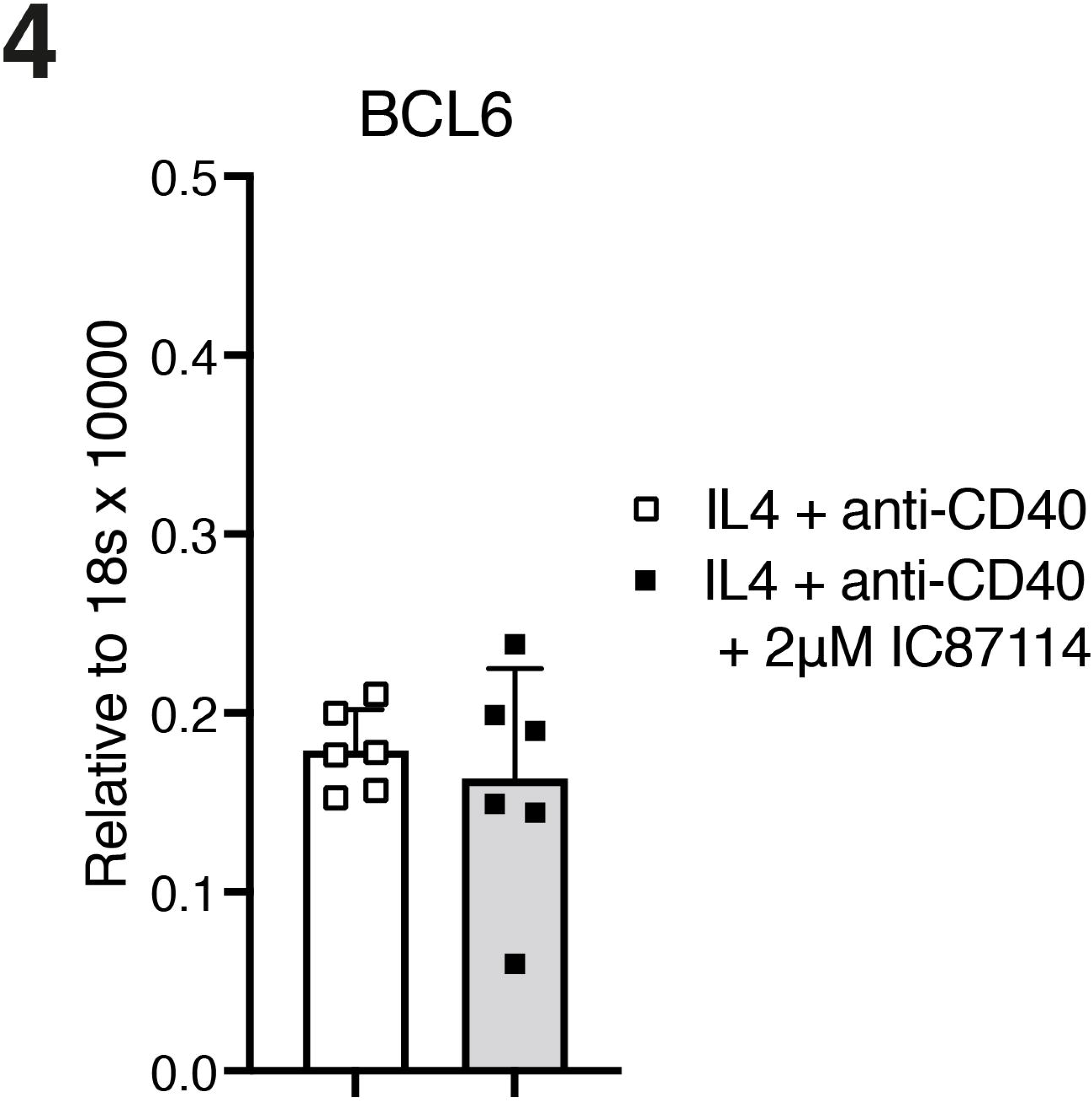
PI3K p110δ inhibition does not affect the expression levels of BCL6 in IL-4 and anti-CD40 stimulated human B cells. The expressions of BCL6 was quantified with qRT-PCR from RNA isolated from day 5 of the tonsillar B cell culture. Gene expression was normalised with 18s rRNA.

**Figure 5.**
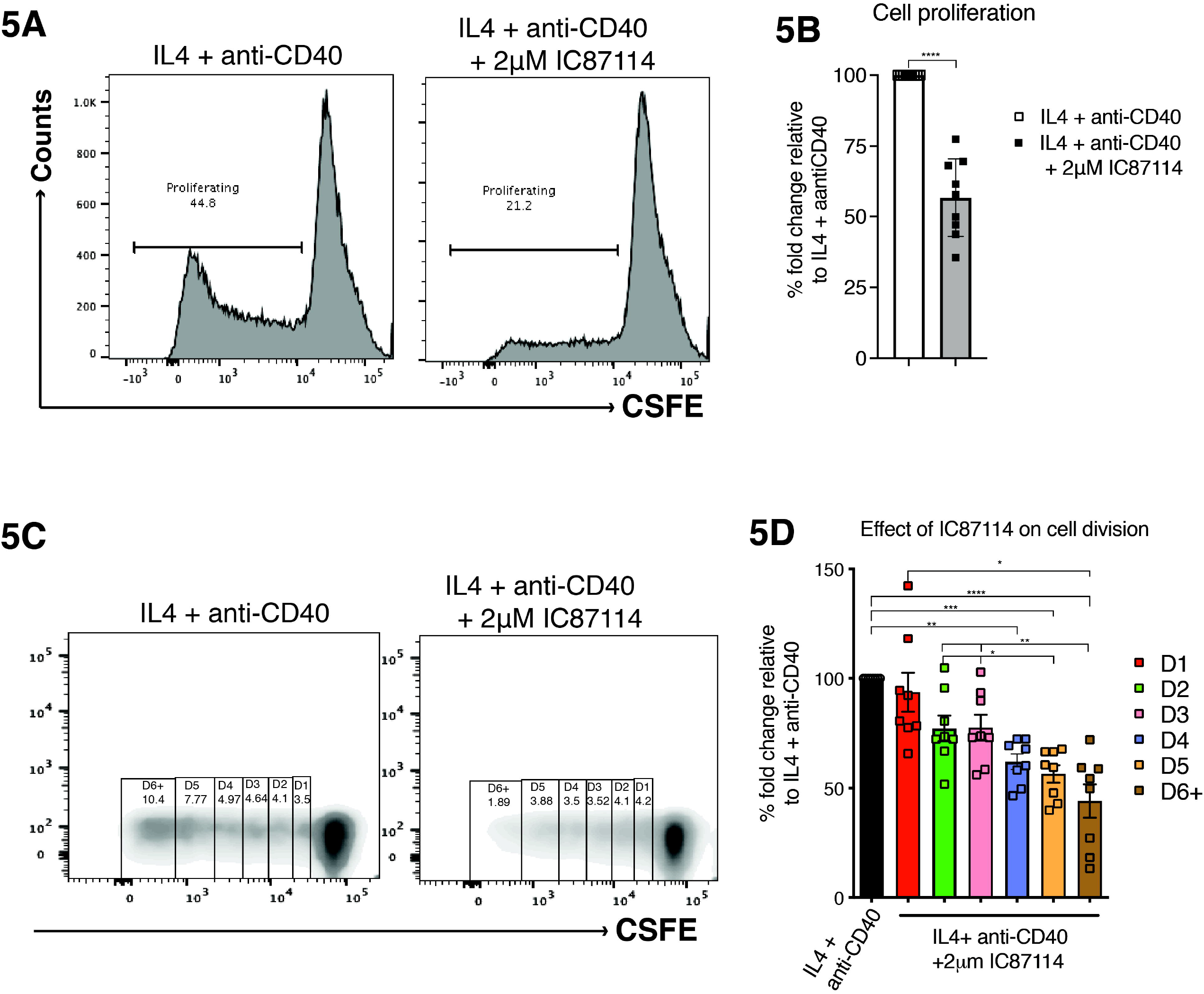
IC87114 inhibits the proliferative capacity of the IL-4 and anti-CD40 stimulated B cells. Isolated tonsil B cells were stained with CFSE and cultured for 10 days with IL-4 and anti-CD40 alone or with 2μM IC87114. **(A)** The FACS histograms show the percentage of proliferating B cells after 10 days of culture, whereas shows the fold change in proliferation relative to the IL-4 and anti-CD40 only culture. The statistical comparisons were performed using the paired t-test. **** P < 0.0001. (**C**) The FACS plots show the number of divisions and the percentage of cells at each division that the IL-4 and anti-CD40 stimulated B cells, with or without IC87114, were undergoing. Based on the CFSE dilution, six numbers of cell divisions (D1-D6+) were identified and gated. (D) Graph shows the frequency of B cells at divisions 1 to 6+ (D1 to D6+).

The data suggests that the modulation of the B cell proliferation is the main mechanism through which PI3K p110δ signalling regulates CSR to IgE in human B cells stimulated with IL-4 and anti-CD40.

## Discussion

Mast cells lacking the p110δ activity have a considerable reduction in IgE-FcεRI mediated degranulation and release of proinflammatory mediators(27,28). Inhibition of the p110δ signalling in mouse models of asthma is also associated with reduced Th2 cytokines as well as reduced lung cellular influx, including tissue esonophilia, airway mucus production and airway hyper-responsiveness(29–31). These studies confirm PI3K p110δ as a potential target for the treatment of allergic disease. Surprisingly, however, other mouse studies have reported that PI3K p110δ signaling negatively regulates CSR to IgE, thus its inhibition enhances IgE production(15–17). In this study, we investigated the effect of PI3K p110δ inhibition in cultures of human tonsil B cells stimulated to undergo CSR to IgE. We find that the inhibition of the PI3K p110δ signaling in these human B cell cultures reduces IgE production. This contradiction between the human and mouse responses to PI3K p110δ inhibition highlights the differences in their B cell regulatory systems of IgE production.

PI3K p110δ is activated downstream of various receptors on B cells and plays critical roles in the development and function of B cells(18,32–34). Several studies have also implicated PI3K signalling in the regulation of CSR(15,17,23,35). The first line of evidence came from mouse B cells lacking PTEN, a phosphatase that negatively regulates PI3K activity(35). These PTEN-deficient mouse B cells, which have enhanced PI3K signalling, fail to undergo CSR both *in vivo* and *in vitro*. IC87114 reverses the effect of PTEN-deficiency on CSR, suggesting that CSR is predominantly regulated by the PI3K p110δ isoform. Furthermore, inhibition of PI3K p110δ was able to also enhance CSR in wild type mouse B cells, suggesting that under normal circumstances PI3K p110δ negatively regulates CSR. In the case of mouse IgE and IgG1, PI3K p110δ negatively regulates their IL4 and anti-CD40 induced CSR *in vitro* by downregulating the levels of AID as well as that of εGLT and γ1GLT, respectively(15,17). Interestingly, our data from the human B cell cultures were inconsistent with these observations made in the mouse systems. IC87114 treatment leads to a significant reduction in the number of IgE^+^ cells in the IL-4 and anti-CD40 stimulated tonsil B cell cultures. The effect of IC87114 on IgG1^+^ cells was not as marked, and a significant reduction was observed only with the highest concentration of 2µM IC87114. It has also been reported that upon immunisation with T dependent antigens, both total and antigen specific IgE titters are dramatically reduced in mice treated with IC87114 and those with B cell deficient PI3K p110δ (CD19^Cre^p110δ^flox/flox^)(15,36). In contrast, the titers of IgG1 and IgM are unaffected, suggesting that the PI3K p110δ selectively regulates IgE production *in vivo*. Further investigations indicated that PI3K p110δ regulates CSR to IgE in a B cell-intrinsic manner by modulating the expression levels of BCL6 (16). However, we did not observe an effect of IC87114 on the class switch signals downstream of IL4R and CD40 and the levels of BCL6 expression were unchanged, suggesting that the mechanisms through which PI3K signalling regulates CSR to IgE differs between the human and the mouse systems.

PI3K p110δ signalling plays a key role in B cell proliferation as it regulates the expression levels of different components of the cell cycle machinery(27,33,34). Our findings show that regulation of the B cell proliferative capacity is the key mechanism through which PI3K p110δ signalling regulates CSR to IgE in our IL4 and anti-CD40 stimulated B cell cultures. Indeed, IC87114 significantly reduces the frequency of cells that have undergone 4 or more cell divisions. This contradicts the previous observations made in mouse, which showed that the proliferation of mouse B cells, stimulated with IL-4 and anti-CD40, is unaffected by the PI3K p110δ inhibition(3,15,17,23). Furthermore, the effect of the IC87114 on the number of B cells undergoing 4 or more cell divisions may explain why we see a more pronounced effect on CSR to IgE, which requires a minimum of 5 divisions, compared to CSR to IgG1, which only requires two cell divisions.

An important feature of both mouse and human IgE+ B cells is their tendency to differentiate into the PC lineage(6,8,9,37). Recent studies in mice revealed that this process is promoted by the mIgE signalling independently of antigen and involves the PI3K signalling pathway(4,5,38). The drastic effects of IC87114 on the amounts of secreted IgE suggests that in addition to inhibiting the CSR to human IgE, the loss of PI3K p110δ signalling might also inhibit the generation of IgE producing PCs. Our future studies will aim to elucidate the role of PI3K p110δ signalling in the differentiation of human IgE+ B cells.

Overall, our data have demonstrated that the regulation of IgE production by PI3K p110δ differs between the mouse and the human systems. In human B cell cultures, the PI3K p110δ regulates IgE production by modulating the proliferative capacity of the IL-4 and anti-CD40 stimulated B cells. In addition, these data highlights the therapeutic potential of small molecule inhibitors of PI3K p110δ in the treatment of allergic disease.

## Abbreviations

AID: Activation-induced cytidine deaminase
CFS: Carboxyfluorescein diacetate succinimidyl ester
CSR: Class switch recombination
FACS: Fluorescence activated cell sorting
GLT: Germline transcription
NFκB: Nuclear factor-kappa B
PI3K: Phosphoinositide 3-kinase
PTEN: Phosphatase and tensin homolog
STAT6: Signal transducer and activator of transcription-6

## Acknowledgments

We are grateful to the patients and the ENT surgical team at the Guy’s & St Thomas’ NHS Foundation Trust for their help and support in the collection of tonsils used in this research.

## Author Contributions

A.C-P., A.d.S, and D.J.F performed experiments and analysed data, and critically reviewed the manuscript. H.J.G. provided guidance, interpreted data and critically reviewed the manuscript. F.R. designed and performed experiments, analysed data and wrote the paper. All authors reviewed the final manuscript.

